# Seasonal Phenology of *Empoasca fabae* (Hemiptera: Cicadellidae) in Québec, Canada

**DOI:** 10.1101/2025.02.21.639576

**Authors:** Abraão Almeida Santos, Fausto Henrique Vieira Araújo, Nicolas Plante, Ricardo Siqueira da Silva, Edel Pérez-Lopéz

## Abstract

Climate change is reshaping insect population dynamics in North America, notably impacting the migratory pest *Empoasca fabae* (Harris) (Hemiptera: Cicadellidae). While its phenology is well studied in the United States, knowledge gaps exist regarding its dynamics in Eastern Canada, one of its northernmost migration areas. Our study integrates degree-day models, CLIMEX ecological niche modeling, and field-collected data from Québec to assess *E. fabae* seasonal phenology and monthly climatic suitability. Our results indicate that *E. fabae* completes one to two generations in Québec, with earlier emergence and higher generational potential in warmer southeastern regions compared to cooler northeastern regions. CLIMEX modeling showed that suitable climatic conditions for *E. fabae* growth begin in April, peak from May to September, and decline by November. First adult captures occurred from late May to early June, with population peaks in June-July and a decrease by September. Observed adult peaks occurred earlier than predicted by degree-day models, suggesting that additional environmental factors, such as wind patterns and host plant availability, influence early-season population dynamics. This study provides a comprehensive understanding of *E. fabae* phenology in Québec and highlights the importance of incorporating climatic and ecological modeling to predict future population trends. Further research on diapause onset, late-season persistence, and migration patterns is needed to refine predictive models and inform pest management strategies in Québec. Understanding these factors will be essential in mitigating potential economic impacts amid ongoing climate change.

## INTRODUCTION

Ongoing climate change is reshaping insect population dynamics in North America, particularly through shifts in overwintering ranges and seasonal phenology, with diverse consequences depending on species and life history (Maredia et al., 1998; Baker et al., 2015; Walter et al., 2018; Lawton et al., 2022). *Empoasca fabae* (Harris) (Hemiptera: Cicadellidae), the potato leafhopper, is a migratory insect pest species native to North America (DeLong, 1938; Ross et al., 1964) whose phenology is likely influenced by these changes. For instance, increased temperatures have led to the earlier arrival of *E. fabae* in northern areas of the United States (USA) and increased crop damage (Maredia et al., 1998; Baker et al., 2015). Although evidence of effects on overwintering populations is growing, it remains unclear how these changes influence *E. fabae* migratory population dynamics in northern areas, particularly in Canada, during the growing season and fall, and how this relates to its baseline phenology.

Each spring, *E. fabae* migrates from its overwintering zone in the southeastern USA to northern areas in the USA and Eastern Canada (DeLong, 1938; Medler, 1957; Specker et al., 1990; Sidumo et al., 2005) with arrival occurring from early May until mid-June, though some early arrivals have been reported by the end of March in North Central USA states (Maredia et al., 1998; Baker et al., 2015). Then, local generations emerge, and population size increases and peaks from June to August (Medler et al., 1966; Sidumo et al., 2005). The population is then expected to decline by September, likely associated with the return of individuals to the overwintering zone, the reduction of suitable hosts, and the beginning of freezing temperatures impacting individual survival and reproduction (Medler et al., 1966; Hogg and Hoffman, 1989; Specker et al., 1990). However, from the end of October to early November, *E. fabae* can still be found outside the overwintering area (Medler et al., 1966).

Québec, Canada, is one of the final northern frontiers where *E. fabae* immigrates every spring (Bostanian et al., 2003; Plante et al., 2024). In this province, studies on *E. fabae* phenology have primarily focused on abundance from mid-May to the end of September (Bostanian et al., 2003; Bostanian et al., 2006; Saguez et al., 2014; Plante et al., 2024). These studies have also aimed to delineate management practices to reduce pest damage in field crops such as alfalfa, often assuming similar life history dynamics (e.g., adult emergence dates and female reproductive diapause) as in northern USA states (Appleton et al., 2003). Valiquette and Comtois (2022) proposed a seasonal model for *E. fabae* in Québec, and this model suggests that the first-generation adult emergence occurs in mid-June and the second by the beginning of August. They also indicated that newborn females enter a reproductive diapause state— associated with the return migration (Taylor et al., 1995) — by mid-August and that non-migratory insects die by October (Valiquette and Comtois, 2022). However, this proposed model did not account for regional variations within the province, and those variations—mainly related to minimum temperature—are suggested to impact the leafhopper population dynamic at regional scales (DeLong, 1938; Hogg and Hoffman, 1989).

Understanding the seasonal phenology of *E. fabae* in Québec and its variations within the province is crucial to estimating further the ongoing changes caused by climate change on this species in Canada (Taylor and Shields, 2018). In this study, we aimed to describe the seasonal phenology of *E. fabae* in four geographical regions of the province using two approaches: (*i*) degree day models based on the thermal thresholds developed for each life stage of *E. fabae* and (*ii*) ecological niche modeling to estimate monthly areas suitability.

## MATERIALS AND METHODS

### Degree days model

We estimate the development time of *E. fabae* based on the accumulated degree days (ADD, °) considering the first day of the year, January 1st, as the null model and the first adult capture on the yellow sticky traps as the first capture model for each region sampled. These two models were performed to estimate the adult emergence date and time and the number of generations based on two scenarios: (*i*) the hypothetical scenario where *E. fabae* is considered a local species considering that relative species from the subtribe Empoascina overwinter in the egg stage, such as *Hebata* (*Empoasca*) *erigeron* (DeLong) (DeLong, 1938) (null model) and (*ii*) the realistic scenario where *E. fabae* migrates to Canada (first capture) (Plante et al., 2024). Daily temperatures for each region (see Field Sampling section) were obtained from the NASA Prediction of Worldwide Energy Resources (POWER) database (version 2.4.3; https://power.larc.nasa.gov/data-access-viewer/) to calculate degree-days (Supplementary Fig. S1). We used this database because no local weather station was available within a 4 km buffer zone (average distance to the nearest station: 7.9 km, range: 4.2–11.84 km). Some nearby stations had incomplete daily or monthly data. To validate the accuracy of the POWER dataset, we performed a Spearman correlation analysis (α = 0.05) comparing the minimum and maximum daily temperatures from the nearest available weather station with the corresponding predicted values for each year and region. Our analysis demonstrated strong agreement between the observed local weather data and the POWER predictions for each year and region (r > 0.97, p < 0.0001; Supplementary Fig. S2).

The temperature thresholds adopted were 9 °C for egg to nymph and 7.6 °C for nymph to adult development (Simonet and Pienkowski, 1980; Hogg, 1985). We calculated the ADD using the degree day package in R (Lyons, 2022) for each stage using the simple average degree days method and the ‘zeroes out’ method when the daily temperature was below the threshold temperature (Mcmaster and Wilhelm, 1997):

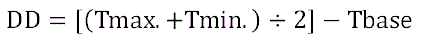

where Tmax is the maximum temperature, Tmin is the minimum temperature, and Tbase is the lower development threshold for each stage. The ADD for each stage was based on the estimations under 18-29 °C (Hogg, 1985) and 13-24 °C (Sher and Shields, 1991). In the warmer regime, the number of degree days accumulated from egg to nymph development is 253°, and from nymph to adult, it is 345° (Σ = 598°) (Hogg, 1985). In the cool regime, those ADDs are 272° and 453°, respectively (Σ = 725°) (Sher and Shields, 1991). Considering the two regimes in both models, we recorded the day as the egg hatch when the ADD reached 253°/272°. From that recorded day, we continued calculating the ADD for nymph development until it reached 345°/453° and noted the day. We recorded this last day as the adult emergence (Σ ADD 598°/725°), and another generation was estimated, assuming an interval of 5 days from the day of adult emergence, as *E. fabae* oviposition initiates on the 6^th^ day after and continues up to 25-50 days after emergence (Sher and Shields, 1991). We conducted these phenological estimations, considering the four fields where sampling was performed (see Field sampling section) in 2021 and 2022. Finally, we averaged both model estimations to determine the first and second generations’ emergence dates (Supplementary Fig. S3 and Supplementary Table S1) following the same approach as Sidumo et al. (2005).

### Climex model

We used the CLIMEX algorithm (version 4.0) to develop spatiotemporal climate suitability for *E. fabae* in Québec (Kriticos et al., 2015). This algorithm generates maps that identify potential areas based on climate conditions favorable for species growth, including predictions on how this suitability varies at temporal scales (e.g., monthly). As a result, it allowed us to identify how suitable areas for *E. fabae* vary over the year in the province.

The CLIMEX algorithm relies on information on species development thermal requirements and a climate database to fit models and generate Ecoclimatic (EI) and Weekly growth (GIw) index maps. The EI results from growth and stress parameters give each species an overall yearly measure of suitable location conditions. EI values range from 0 to 100, where 0 means the area is inadequate for species establishment; 0–30 means that the area has less favorable climate conditions for species growth and development; and > 30 means that the area has highly favorable climate conditions. On the other hand, the GIw index indicates favorable climate conditions over a week on a scale from 0 to 1, where values close to 0 indicate inappropriate conditions for species growth. The conditions are more favorable as GIw values are closer to one (Kriticos et al., 2015).

We used the information in the literature to establish the temperature thresholds for *E. fabae* development and diapause. At the same time, humidity indices and adjusted values were based on the areas of known occurrence of the species. For the EI model, we used the CliMond 101 grid climatic data (https://www.climond.org/) at a spatial resolution of 10 km. The climatic data represent long-term values, including monthly averages of total precipitation (Ptotal), maximum (Tmax), and minimum (Tmin) temperatures, as well as relative humidity at 09:00 (RH 09:00) and 15:00 (RH 15:00) for recent historical climate (Kriticos et al., 2012). For the GIw model, a monthly time series from the Climate Research Unit (version 3.26, https://crudata.uea.ac.uk/cru/data/hrg/), covering the period from 1901 to 2019, was used at also 10 km of spatial resolution.

### Climex model calibration

#### Temperature index

We set the following temperature parameters: lower temperature threshold (DV0) as –9 °C (Decker and Cunningham, 1967; Sidumo et al., 2005), lower optimal temperature (DV1) as 13 °C, the upper optimal temperature (DV2) as 31 °C, and the upper-temperature threshold (DV3) as 38 °C (Kouskolekas and Decker, 1966) (Table 1).

**Table 1.** Final index values to model the seasonal climate suitability of *Empoasca fabae* based on the CLIMEX algorithm. ^#^ soil moisture indices are dimensionless (0 = over dry, 1 = indicates that soil moisture content is 100% capacity).

| Index | Parameter | Value |
| --- | --- | --- |
| Temperature | DV0 = lower threshold | -9 °C |
|  | DV1 = lower optimum temperature | 13 °C |
|  | DV2 = upper optimum temperature | 31 °C |
|  | DV3 = upper threshold | 38 °C |
| Moisture | SM0 = lower soil moisture threshold | 0.1<sup>#</sup> |
|  | SM1 = lower optimum soil moisture | 0.4<sup>#</sup> |
|  | SM2 = upper optimum soil moisture | 1.0<sup>#</sup> |
|  | SM3 = upper soil moisture threshold | 1.5<sup>#</sup> |
| Diapause | DPD0 = diapause induction day length | 10 hours |
|  | DPT0 = diapause induction temperature | -9 °C |
| Cold Stress | TTCS = cold stress temperature threshold | -9 °C |
|  | THCS = stress accumulation rate | -0.00003/week |
| Heat Stress | TTHS = heat stress temperature threshold | 38 °C |
|  | THHS = stress accumulation rate | 0.0001/week |
| Wet Stress | SMWS = wet stress threshold | 1.5<sup>#</sup> |
|  | HWS = stress accumulation rate | 0.002/week |
| Dry Stress | SMDS = dry stress threshold | 0.1<sup>#</sup> |
|  | HDS = stress accumulation rate | -0.007/week |
| Degree Days | PDD = degree days per generation | 725° |

#### Moisture index

We defined the lower soil moisture threshold (SM0) as 0.1 and the lower ideal soil moisture (SM1) as 0.4 to be consistent with the soil moisture of those regions with high *E. fabae* occurrence records. The upper ideal soil moisture (SM2) was set at 1, indicating that the soil moisture content is 100% of its capacity. In contrast, the upper soil moisture threshold (SM3) was set at 1.5, representing areas with excessive soil moisture (Kriticos et al., 2015).

#### Diapause index

*E. fabae* reproductive diapause is associated with photoperiod conditions ≤ 10 hours at both high or low temperatures (Taylor et al., 1995). In addition, Sidumo et al., (2005) estimated the overwintering temperature range for *E. fabae* at –9 °C. Thus, we defined the diapause induction day length (DPD0) as 10 hours and the diapause induction temperature (DPT0) as –9 °C. *Cold and heat stress* The cold stress temperature threshold (TTCS) was set at –9 °C (Sidumo et al., 2005), and the stress accumulation rate (THCS) was set at –0.00003 week^-1^. These values were adjusted to suit the cold regions of Canada (Kriticos et al., 2015). We set the heat stress temperature threshold (TTHS) at 38 °C (Kouskolekas and Decker, 1966), and the stress accumulation rate (THHS) was 0.0001 week^-1^ (Kriticos et al., 2015).

#### Wet and dry stress

The wet stress threshold (SMWS) was set at 1.5, while the stress accumulation rate (HWS) was defined as 0.002 week^-1^. The dry stress threshold (SMDS) was fixed at 0.1, with the stress accumulation rate (HDS) set at –0.007 week^-1^. These values were adjusted for *E. fabae* to achieve a high Ecoclimatic Index (EI) in the USA and Canada.

#### Degree days

As previously indicated, two-degree days models were determined for *E. fabae* (Hogg, 1985; Sher and Shields, 1991). The final model used as the degree-days per generation (PDD) was 725° because it provided a more accurate model consistent with the known distribution of *E. fabae*.

### Climex model validation

We validated the model based on the correspondence between EI-suitable categories (i.e., > 0) and the occurrence records of *E. fabae*. We compiled the occurrence of *E. fabae* from the Global Biodiversity Information Facility website (GBIF, 2024). A total of 7,459 records were distributed across North America and then plotted in the final ecoclimatic index map.

For the GIw model, we performed a Spearman correlation (α = 0.05) between the number of adults caught (see Field sampling section) and the GIw for the corresponding week. We extracted the GIw for each sampled field from 2010 to 2019 and then used the averaged values to conduct the correlation analysis (Supplementary Table S2). We used this approach because monthly climate data for the field sampling years (2021–2022) were unavailable.

### Field sampling

We used a leafhopper collection survey conducted on strawberry farms in Québec during the growing seasons of 2021 and 2022 (Plante et al., 2024) to depict *E. fabae* seasonal phenology, comparing it with the degree days models and CLIMEX predictions. We selected one farm per geographical region explored by Plante et al. (2024), making a total of 4 farms with distances between farms ranging from 38 to 210 km (Fig. 1a-b). Briefly, each year, from the third week of May until the end of September, surveys were conducted using yellow sticky traps attached to wood pins placed at the level of the strawberry plants and changed weekly (Fig. 1c). The identification of *E. fabae* was based on external morphology (Fig. 1d) and male genitalia dissection using the key developed by Beirne (1956). For confirmation, we compared the identified specimens with those from the Canadian National Collection of Insects, Arachnids, and Nematodes in Ottawa (Ontario, Canada). Since the traps were not monitored daily after the first installation week for logistical reasons, we adopted a conservative approach, considering the first capture day as the last one when the trap was in the field when the first *E. fabae* was captured in each region (Table 2).

**Fig. 1.**
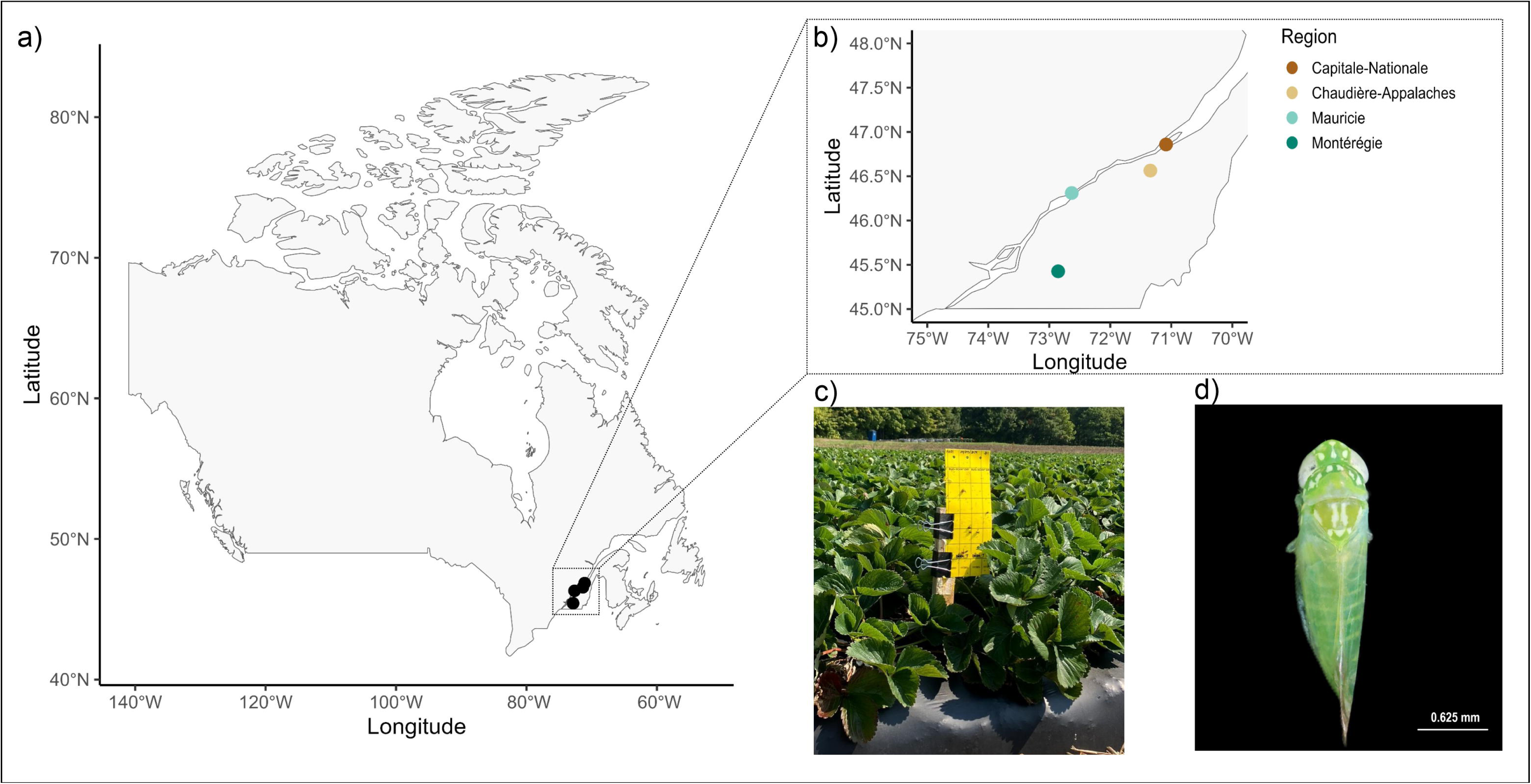
Leafhoppers sampling information. (a-b) The geographical position of fields in Québec (Canada). (c) Yellow stick trap used to capture *Empoasca fabae* in the strawberry fields. (d) *Empoasca fabae* adults with the primary morphological feature for identification based on the six white spots on the top of the head behind the eyes.

**Table 2.**
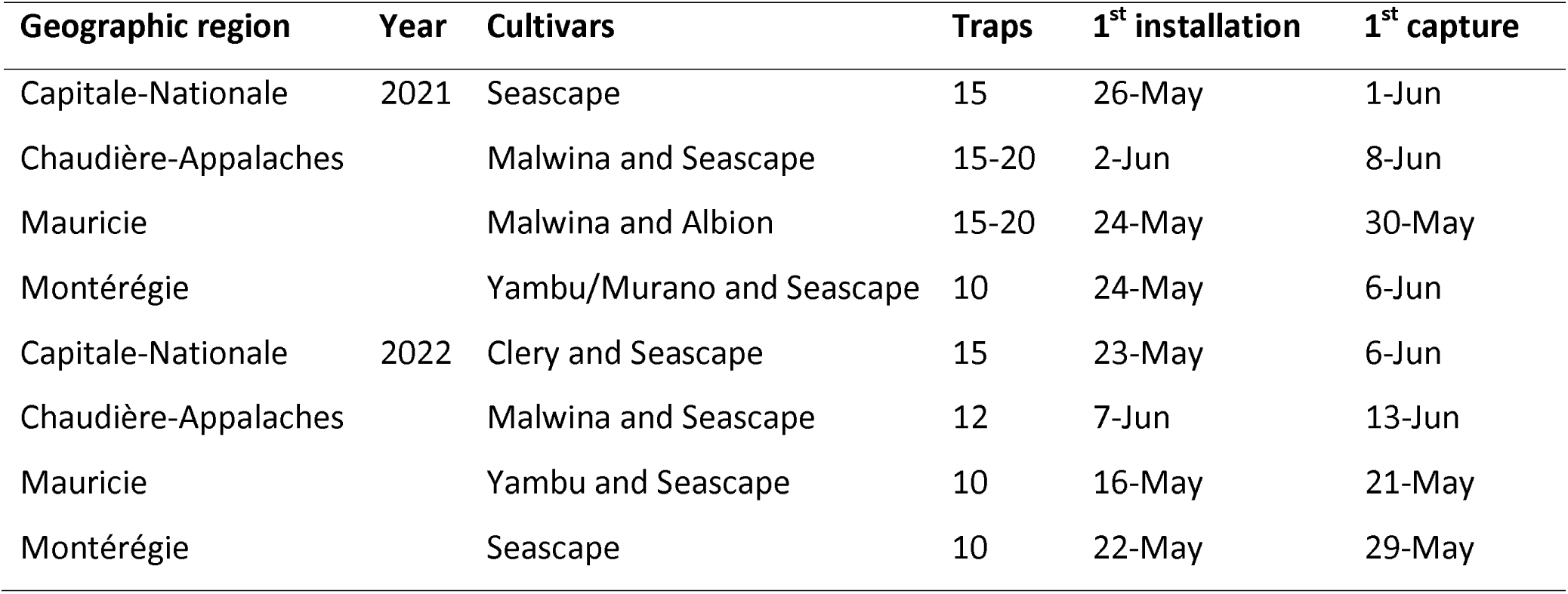
Sampling areas information related to each geographic region, year, the strawberries cultivars, number of traps per week, first installation and capture of *Empoasca fabae* in Québec, Canada.

## RESULTS

### Estimating adult’s emergence

By combining the Hogg (1985), the Sher and Shields (1991) degree days estimations, and the null and first capture models, we found that the date of the first-generation adult emergence of *E. fabae* in Québec ranged (minimum and maximum) from 9 July to 18 August, while the second from 10 September to 25 October (Table 3). Interestingly, these models indicated that southeast regions of the province (Mauricie and Montérégie) had early emergence dates compared to northeast ones (Capitale-Nationale and Chaudière-Appalaches). Nevertheless, the first capture model suggests that a second generation of insects would be expected only in Mauricie and Montérégie (Table 3). The number of days required for the first adult emergence was longer (90-124 days) than the second (47-90 days) in the null model. In the first capture models, the first (53-79 days) and second (49-85 days) generations took similar emergence times (Table 3).

**Table 3.** Estimated date and number of days for *Empoasca fabae* adult emergence (first and second generation) in four geographic regions in Québec, Canada. The null model considers *E. fabae* as an overwintering species in Canada, while the first capture is based on the first date of adult capture in yellow stick traps. The values are averaged according to the accumulated degree-days models of Hogg (1985) and Sher and Shields (1991).

| Geographic region | Year | Emergence (null model) |  | Emergence (first capture) |  |
| --- | --- | --- | --- | --- | --- |
|  |  | F1 | F2 | F1 | F2 |
| Capitale-Nationale | 2021 | 06 Aug, 120 d | 22 Oct, 77 d | 14 Aug, 75 d | - |
|  | 2022 | 07 Aug, 97 d | - | 18 Aug, 79 d | - |
| Chaudière-Appalaches | 2021 | 26 Jul, 124 d | 23 Sep, 60 d | 12 Aug, 66 d | - |
|  | 2022 | 27 Jul, 124 d | 25 Oct, 90 d | 16 Aug, 65 d | - |
| Mauricie | 2021 | 21 Jul, 119 d | 25 Sep, 62 d | 01 Aug, 64 d | 23 Sep, 53 d |
|  | 2022 | 23 Jul, 90 d | 21 Sep, 59 d | 30 Jul, 71 d | 23 Oct, 85 d |
| Montérégie | 2021 | 09 Jul, 121 d | 31 Aug, 47 d | 28 Jul, 53 d | 21 Sep, 50 d |
|  | 2022 | 13 Jul, 119 d | 10 Sep, 54 d | 28 Jul, 61 d | 16 Sep, 49 d |

### Climex models

The CLIMEX model performed well with the final set of parameters for *E. fabae* (Table 1). When this model was validated with the occurrence records of *E. fabae*, all occurrences aligned with moderate and high suitability areas (EI > 0; Supplementary Fig. S4), then, we could estimate how this suitability would vary monthly using the seasonal model (GIw).

The seasonal model showed that the suitability of Québec for *E. fabae* varies monthly with periods of unsuitable and high conditions for species growth (Fig. 2). From January to March, the GIw values were closer to zero. Then, GIw increased in April, with the highest values from June to August (0.5-0.6), subsequently decreasing. The GIw values for the four regions showed a similar pattern, with the highest values from May to September and decreasing afterward (Fig. 3). These values also presented a high variability (standard deviation) based on the period evaluated from 2010 to 2019 (Fig. 3). No significant correlation was found between the number of adults and GIw values (r = –0.006, p = 0.94).

**Fig. 2.**
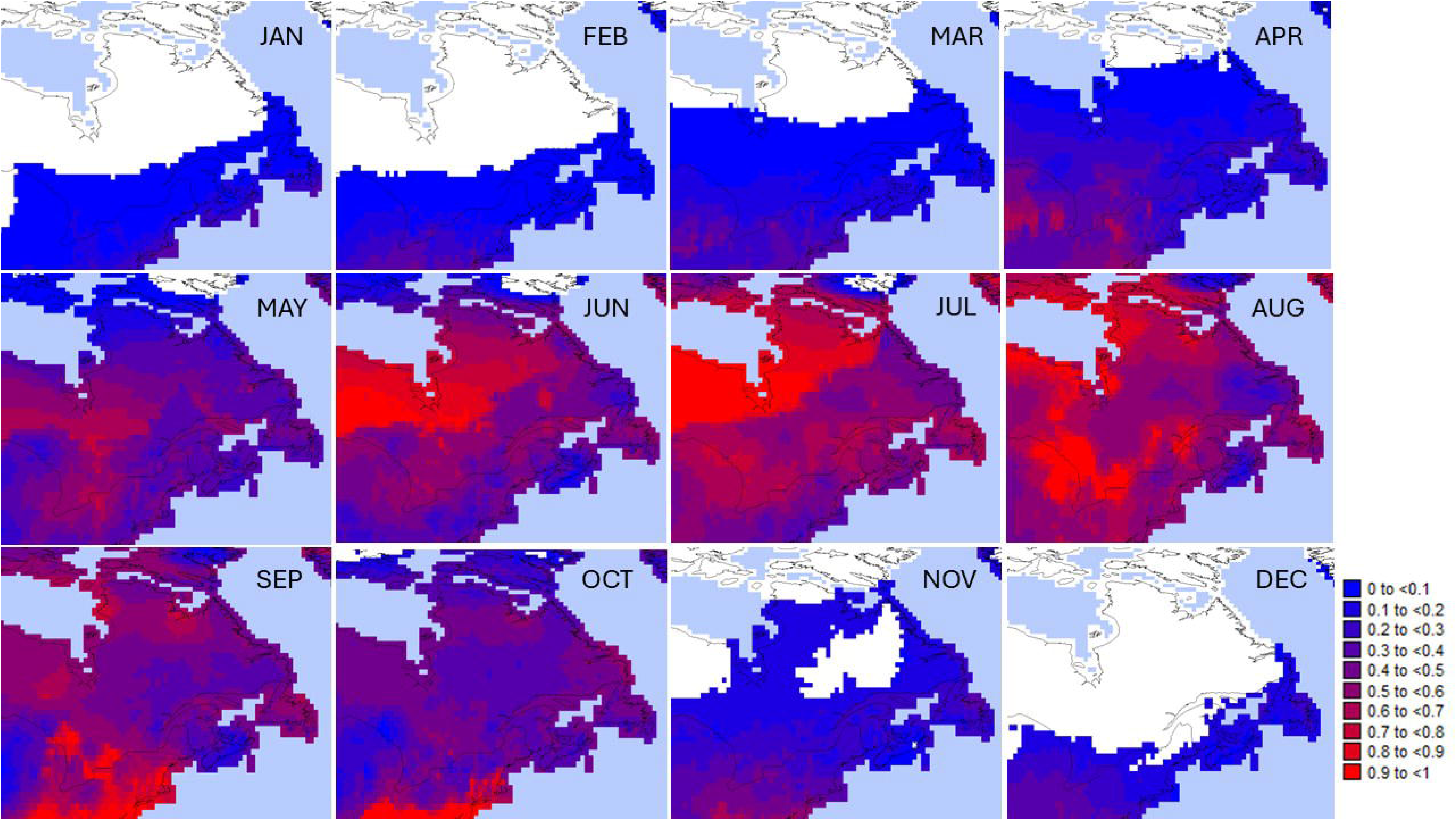
Seasonal climate suitability based on the weekly Growth Index (GIW) for *Empoasca fabae* in Québec (Canada). The index varied from 0 to 1, where values > 0.1 indicate growing conditions for the species.

**Fig. 3.**
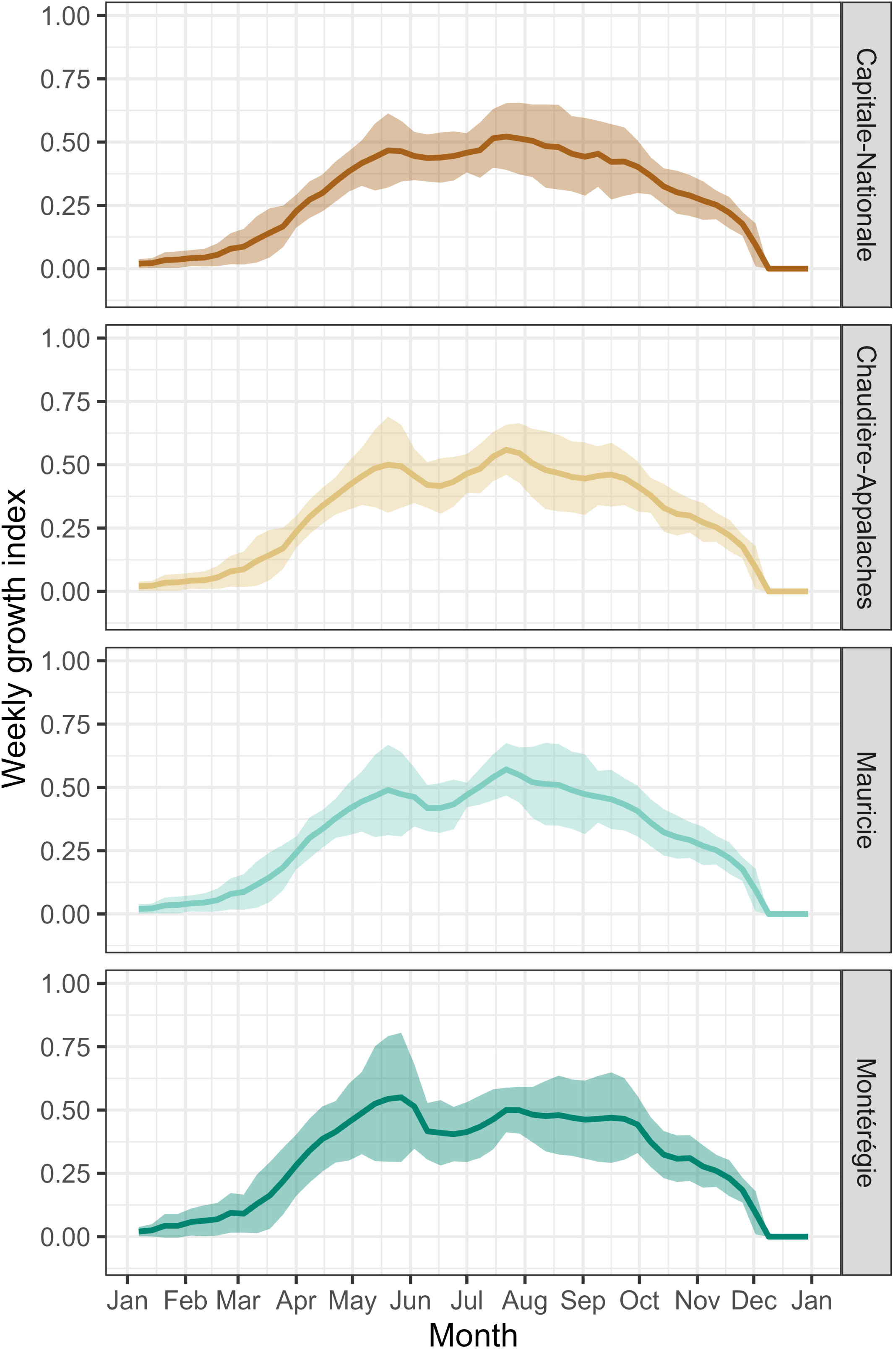
Estimated weekly growth index based on the seasonal model using the CLIMEX algorithm for each geographical region in Québec, Canada. The line indicates the average and shadows ± standard deviation from 2010 to 2019. The index varied from 0 to 1, where values > 0.1 indicate growing conditions for the species.

### Field sampling

The first captures of *E. fabae* occurred at the end of May and the beginning of June (Table 2). Interestingly, the peak of adults captured varied among regions from the end of June to the middle of July. In all cases, the number of adults captured was lower by September, with no adults detected in the yellow sticky traps by the end of that month (Fig. 4). When we compared the peaks of adults captured with the estimated first emergence dates by the two models (null and first capture), the peak was before the predicted dates in both cases (Fig. 4), except in Chaudière-Appalaches (2021) and Mauricie (2022). Considering the GIw values, the first capture occurred only by May when values were > 0.40.

**Fig. 4.**
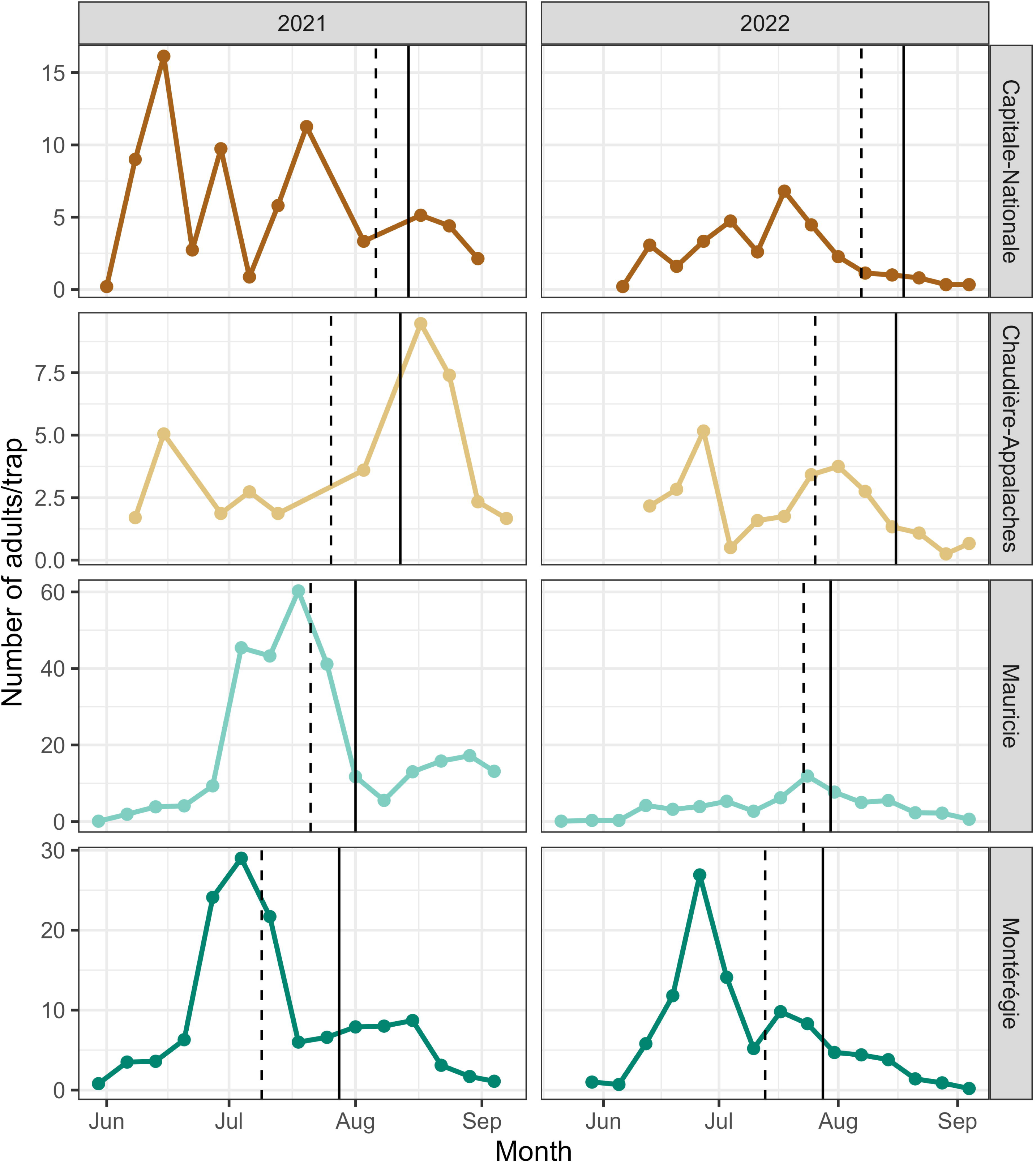
Average number of *Empoasca fabae* adults collected per trap in each Québec, Canada geographical region. The dotted and solid lines indicate the first adult emergence based on the degree days null and first capture models.

## DISCUSSION

Québec is one of the northernmost regions to which *E. fabae* migrates annually from its overwintering zone in the Southeastern USA (Sidumo et al., 2005). While numerous studies have investigated the phenology and population dynamic of this species in the USA (e.g., Delong, 1928; DeLong, 1931; Poss, 1932; Delong and Caldwell, 1935; DeLong, 1938; Medler, 1957; Pienkowski and Medler, 1964; Kouskolekas and Decker, 1966; Medler et al., 1966; Simonet and Pienkowski, 1980; Taylor and Reling, 1986; Lamp et al., 1989; Parr and Pass, 1989; Carlson et al., 1992; Taylor et al., 1995; Taylor and Shields, 1995; Maredia et al., 1998; Sidumo et al., 2005; Baker et al., 2015; Lagos-Kutz et al., 2024), research in Canada has primarily focused on economic aspects related to pest control in field crops (e.g., Cheng and Roy, 1985; Appleton et al., 2003; Shi, 2023). In Québec, research has emphasized the phenology of *E. fabae* from May to September (Bostanian et al., 2003; Bostanian et al., 2006; Saguez et al., 2014; Plante et al., 2024), with limited information available for the earlier and later parts of the year, and a seasonal model for the province proposed (Valiquette and Comtois, 2022). By combining degree-day models and a monthly ecological model using the CLIMEX algorithm, we gained insights into the annual phenology of *E. fabae* in Québec, which will contribute to a better understanding of its population dynamics.

Across North America, *E. fabae* generations typically range from one to two, with a partial third-fourth generation possible (DeLong, 1938; Sidumo et al., 2005). Our findings suggest the potential for two *E. fabae* generations in Québec, aligning with the trend of decreasing generation numbers with increasing latitude across North America (Sidumo et al., 2005). However, the first capture model revealed regional variations within the province, with southeast regions showing a higher likelihood of supporting two generations. In contrast, northeast regions may only support a single generation.

While humidity and precipitation are generally considered primary factors influencing *E. fabae* occurrence, it has been suggested that under stable wet conditions, temperature becomes the dominant factor influencing development time and population size (DeLong, 1938; Baker et al., 2015). This scenario appears relevant in Québec, as previous reports indicated a lack of significant association between rainfall and Nearctic leafhopper population size (Plante et al., 2024). Conversely, temperature, particularly minimum temperatures during the growing season, likely plays a crucial role in accelerating development, potentially enabling the completion of two generations in warmer regions while limiting development to a single generation in cooler regions (DeLong, 1938; Medler et al., 1966).

Earlier adult emergences observed in the southeast regions align with warmer growing season temperatures, whereas lower temperatures are more common in the northeastern regions (Supplementary Fig. S1). Cold conditions extend *E. fabae* development time (Hogg, 1985; Sher and Shields, 1991), inhibit oviposition between 10-21°C (Taylor et al., 1995) and below 15.6°C (Kieckhefer and Medler, 1964), and increase mortality in immature stages (Simonet and Pienkowski, 1980; Specker et al., 1990). Mortality in adults induced by freezing conditions begins at –9 °C and becomes complete between –13°C and –14°C (Decker and Maddox, 1967). Consequently, colder conditions in the northeast may extend *E. fabae* development time, delay the onset of reproduction, and increase mortality rates, ultimately limiting the number of generations completed within the growing season (DeLong, 1938; Medler et al., 1966). Furthermore, local microclimatic variations within each region, such as those associated with elevation, vegetation cover, and proximity to water bodies, can further influence temperature and contribute to the observed regional differences in *E. fabae* phenology. These findings suggest that environmental heterogeneity, primarily driven by minimum temperature variations across the province, could primarily explain the observed regional differences in *E. fabae* phenology in Québec.

Our seasonal model indicates that suitable conditions (GIw > 0.1) for *E. fabae* development appear in April, with the highest values from May to September (0.4-0.6), aligning with the phenology described during this time frame in the growing season in Québec. In vineyards, the first collections of adults typically occur at the beginning of June, with a peak in July and August, while nymphs appear in June, peaking in July and August, and are not found at the end of September (Bostanian et al., 2003). The first collection in alfalfa fields is similar to that in vineyards but shows multiple population peaks in July in southeastern Québec, whereas only one peak occurs in the northeast (Shi, 2023). Our field sampling also observed an early adult peak from June to July, which differs from the later peak in alfalfa and vineyards, suggesting potential differences in *E. fabae* population dynamics based on host plant availability.

The first capture in our field sampling occurred between late May and early June, coinciding with a GIw value exceeding 0.40, which only appears in May. This aligns with historical records of *E. fabae* first captures in the Northeastern USA (e.g., New York), typically recorded from mid-May to early June, with no reports in April (Medler, 1957; Maredia et al., 1998; Baker et al., 2015). These findings suggest that the onset of favorable spring conditions likely constrains the arrival of migratory individuals in Québec. Notably, none of our models fully capture the complex factors influencing arrival and establishment timing, such as continental winds in the overwintering zone and host plant phenology.

Although our field data collection ended in late September, our model indicated a significant decline in suitable conditions for *E. fabae* by November. In new ongoing sampling efforts, we are extending the sampling period in strawberry fields to November, which will allow us to determine the latest presence of adult *E. fabae*. This effort corresponds with model predictions of suitable conditions in October, highlighting the importance of extended sampling for a more complete understanding of *E. fabae* phenology in Québec.

Our models primarily focused on temperature and precipitation to describe the population dynamics of *E. fabae* in Québec. However, it is crucial to acknowledge that other factors significantly influence populations. These factors include: (a) continental winds, which are associated with the arrival of migratory individuals and the dispersal of populations (Taylor and Reling, 1986; Taylor and Shields, 2018); (b) polyphagy, as the ability of *E. fabae* to utilize a wide range of host plants impacts their survivorship and fecundity (Lamp et al., 1994); (c) natural enemies, which can exert top-down control on *E. fabae* populations, although to a minor extent (Hogg and Hoffman, 1989); and (d) the high degree of generation overlap, which can lead to complex interactions within and between generations due to females’ long reproductive lives, potentially contributing to population outbreaks (Hogg and Hoffman, 1989). These factors were not incorporated into our models, highlighting the need for future research to investigate their interactive effects on *E. fabae* populations in Québec.

Our study aimed to describe the seasonal phenology of *E. fabae* in Québec, providing a data-driven assessment of its population dynamics. While some findings support Valiquette and Comtois’s (2022) proposed model, some discrepancies emerged. Our estimates indicate that the first-generation emergence occurs in July rather than June, aligning even with the null model (i.e., *E. fabae* as a local species), which predicts the first-generation emergence by July. However, our models agree with Valiquette and Comtois (2022) regarding the second generation’s emergence in August-September. Notably, while their model suggests adult *E. fabae* mortality by mid-to-late October, our findings indicate that unfavorable conditions dominate only in November. Given *E. fabae*’s cold hardiness, temperatures below –9 °C—expected to cause complete mortality (Decker & Maddox, 1967; Sidumo et al., 2005)—are typically reached in November rather than October (Supplementary Fig. S1). Although field crops may no longer be available at that time, *E. fabae* is polyphagous and known to feed on pine species (Lamp et al., 1994), which remain present year-round and may serve as late-season hosts, as observed in the overwintering zone (Taylor and Shields, 1995; Sidumo et al., 2005).

The initiation of reproductive diapause in *E. fabae* in Québec remains an open question. Valiquette and Comtois’s (2022) proposed model suggests diapause onset (i.e., newly emerged females in a complete diapause state) in the second week of August. However, field studies in New York (1991 to 1992) indicated that females enter a complete reproductive diapause in late August to early September (Taylor et al., 1995). Temperature data from our study provide some support for this later timing, as conditions within the temperature range of 10-21 °C, known to inhibit oviposition even under photoperiods exceeding 15 hours per day (Taylor et al., 1995), occur more frequently in late August (Supplementary Fig. S1). Furthermore, photoperiods below 14 hours per day are associated with diapause induction (Taylor et al., 1995), begin in late August (Supplementary Fig. S5). Thus, females are more likely to enter full reproductive diapause in late August rather than earlier. However, determining the precise timing of diapause onset requires detailed dissections of females throughout the season to assess their reproductive status.

Rising temperatures are reshaping *E. fabae* phenology, leading to earlier arrivals and increased crop damage in the USA (Maredia et al., 1998; Baker et al., 2015). While climate models predict a northward range expansion of *E. fabae* and other leafhopper pests in North America (Santos et al., 2024), our current understanding of its seasonal phenology in Québec remains incomplete. Key questions remain: (a) What is *E. fabae* spatial distribution within Québec following their arrival? (b) Does it reach northern areas, including non-agricultural habitats in the Québec subarctic and arctic? (c) When the adult population peaks, does it indicate multiple migratory events or local population buildup? (d) When does female reproductive diapause begin? And perhaps most importantly, (e) What happens to *E. fabae* populations in Québec when environmental conditions become unsuitable?

## AUTHOR CONTRIBUTIONS

**Abraão Almeida Santos**: Conceptualization, Data curation, Investigation, Formal analysis, Methodology, Visualization, Validation, Supervision, Software, Writing – original draft, Writing – review & editing. **Fausto Henrique Vieira Araújo**: Investigation, Formal analysis, Visualization, Software, Writing original draft. **Nicolas Plante**: Conceptualization, Data curation, Investigation, Methodology, Validation, Writing – review & editing. **Ricardo Siqueira da Silva**: Supervision, Writing – review & editing. **Edel Pérez-Lopez**: Conceptualization, Funding acquisition, Project administration, Resources, Supervision, Writing – review & editing.

## ACKNOWLEDGMENTS

Fausto Henrique Vieira Araújo and Ricardo Siqueira da Silva acknowledge the following agencies for general support: Conselho Nacional de Desenvolvimento Científico e Tecnológico (CNPq), the Fundação de Amparo à Pesquisa do Estado de Minas Gerais (FAPEMIG) and the Coordenação de Aperfeiçoamento de Pessoal de Nível Superior – Brasil (CAPES) – Finance Code 001.

