## Supplementary Figures for "Seasonal Phenology of *Empoasca fabae* (Hemiptera: Cicadellidae) in Québec, Canada"

**Supplementary Table S1.** Accumulated degree days models based on the estimations of Hogg (1985) and Sher and Shields (1991), and the null and first capture model of *Empoasca fabae* in four geographic regions in Québec, Canada. The null model considers *E. fabae* as an overwintering species in Canada, while the first capture is based on the first date of adult capture in yellow stick traps.

**Supplementary Table S2.** Estimated weekly growth index based on the seasonal model using the CLIMEX algorithm for each geographical region in Québec, Canada, from 2010 to 2019. The index varied from 0 to 1, where values > 0.1 indicate growing conditions for the species.

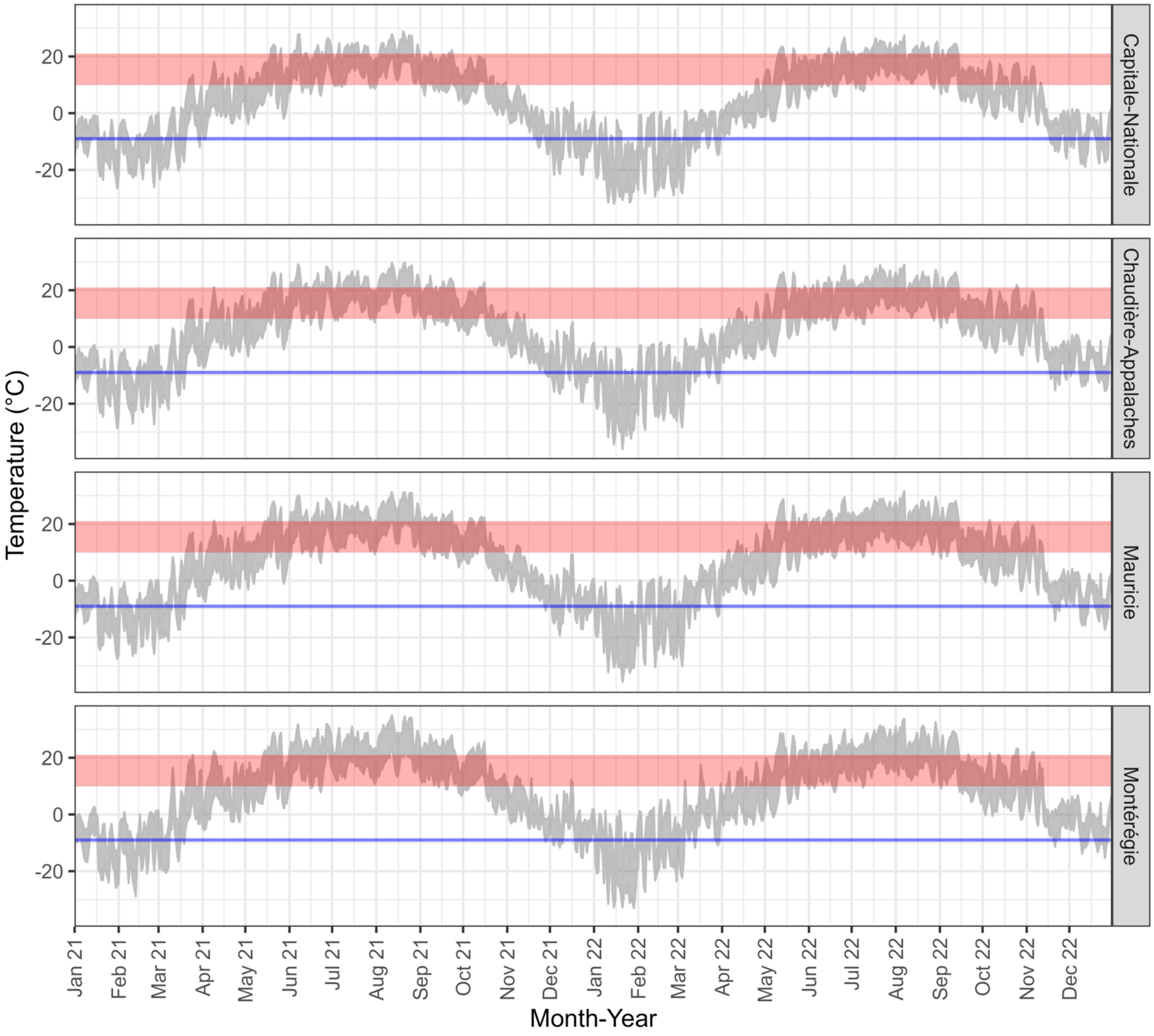


**Supplementary Figure S1.** Temperature (minimum and maximum, °C) in the four geographic regions of Québec (Canada). The red rectangle indicates the temperature window (10-21 °C) where the oviposition of *Empoasca fabae* is inhibited under laboratory conditions. The blue line indicates the survival temperature threshold for the species in its overwintering zone (-9 °C; Decker and Maddox, 1967; Sidumo et al., 2005).

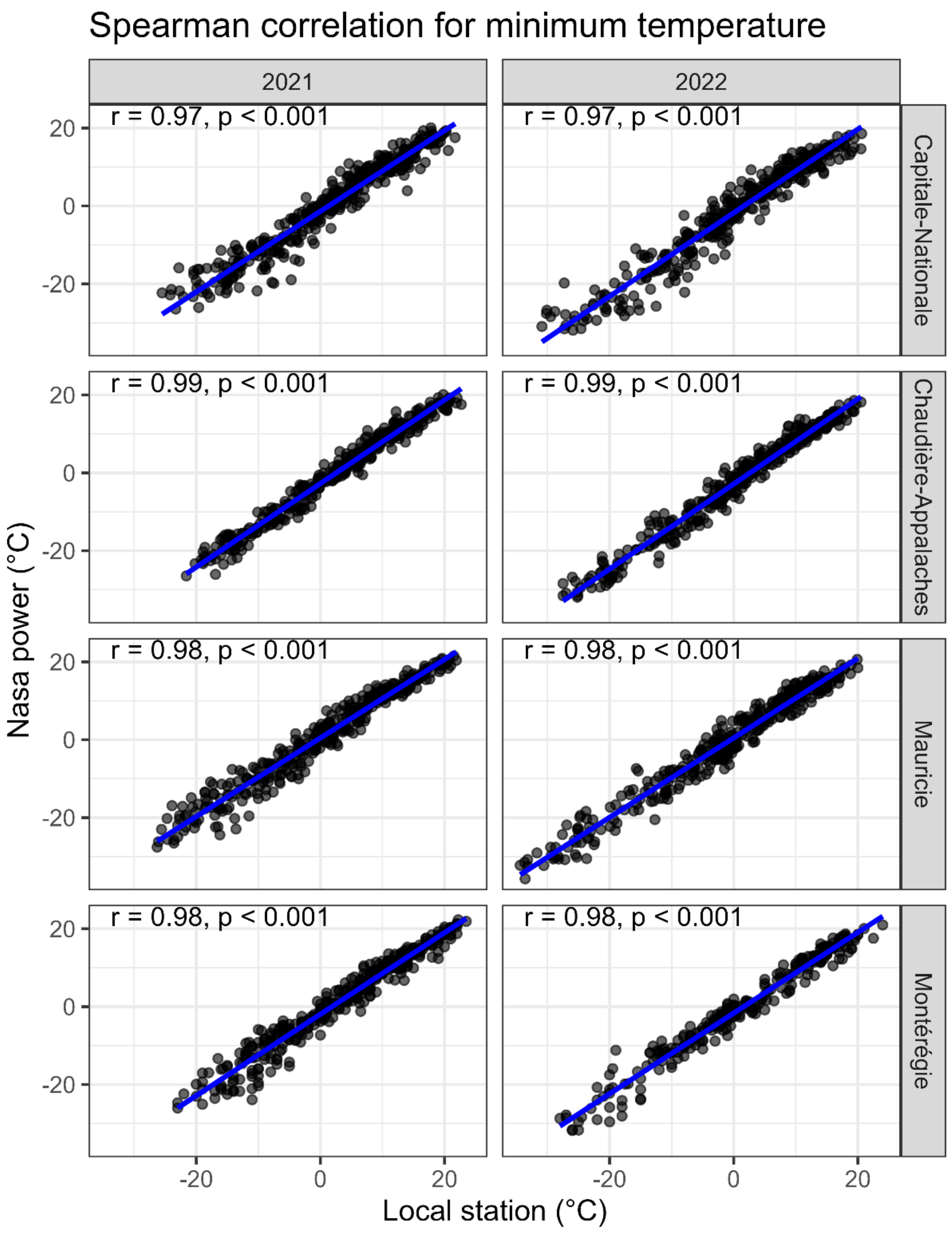


**Supplementary Figure S2(A).** Correlation values for minimum temperature between the closest local weather stations and NASA power prediction for the same site as the weather station.

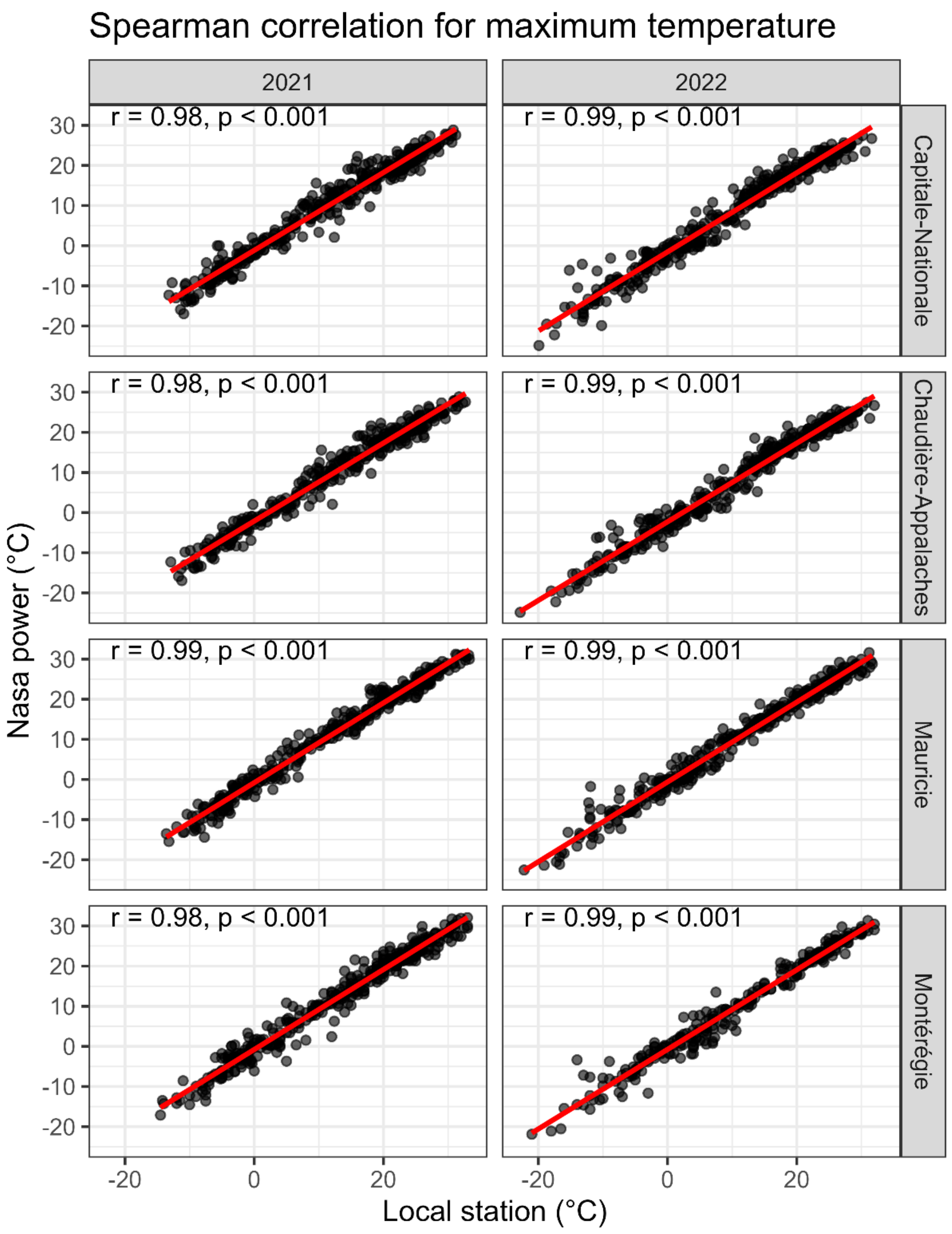


**Supplementary Figure S2(B).** Correlation values for maximum temperature between the closest local weather stations and NASA power prediction for the same site as the weather station.

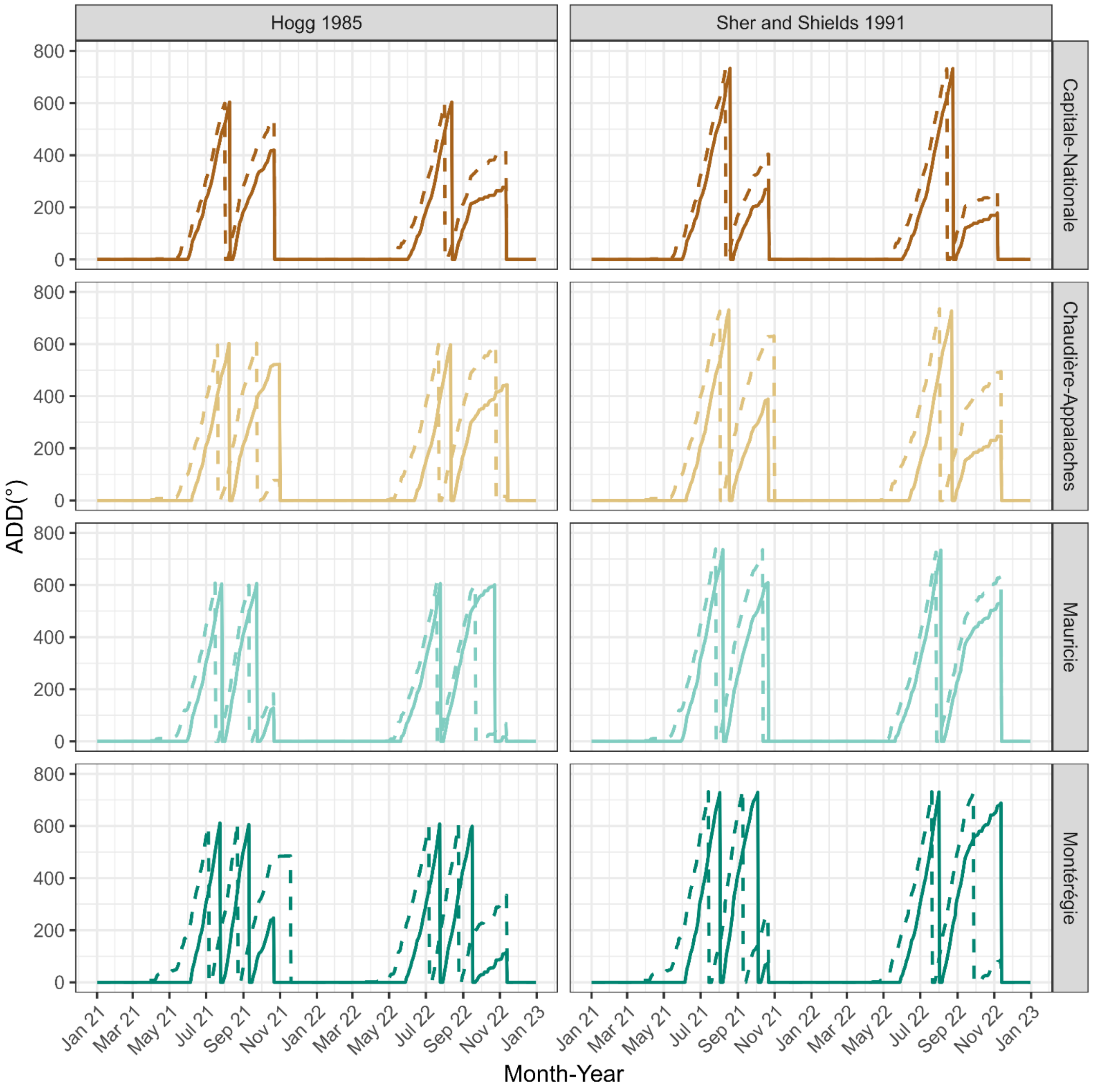


**Supplementary Figure S3.** The accumulated degree days models (ADD, °) for *Empoasca fabae* development from egg to adult based on Hogg (1985) (Σ = 598°, 18-29 °C) and Sher and Shields (1991) (Σ = 726°, 13-24 °C ) for each geographic region according to a null and first capture. The null model (dashed line) considers the ADD since January 1$^{st}$ (i.e., species overwinter in Québec, Canada), while the first capture (solid line) considers the date when the adults were captured using the yellow stick traps.

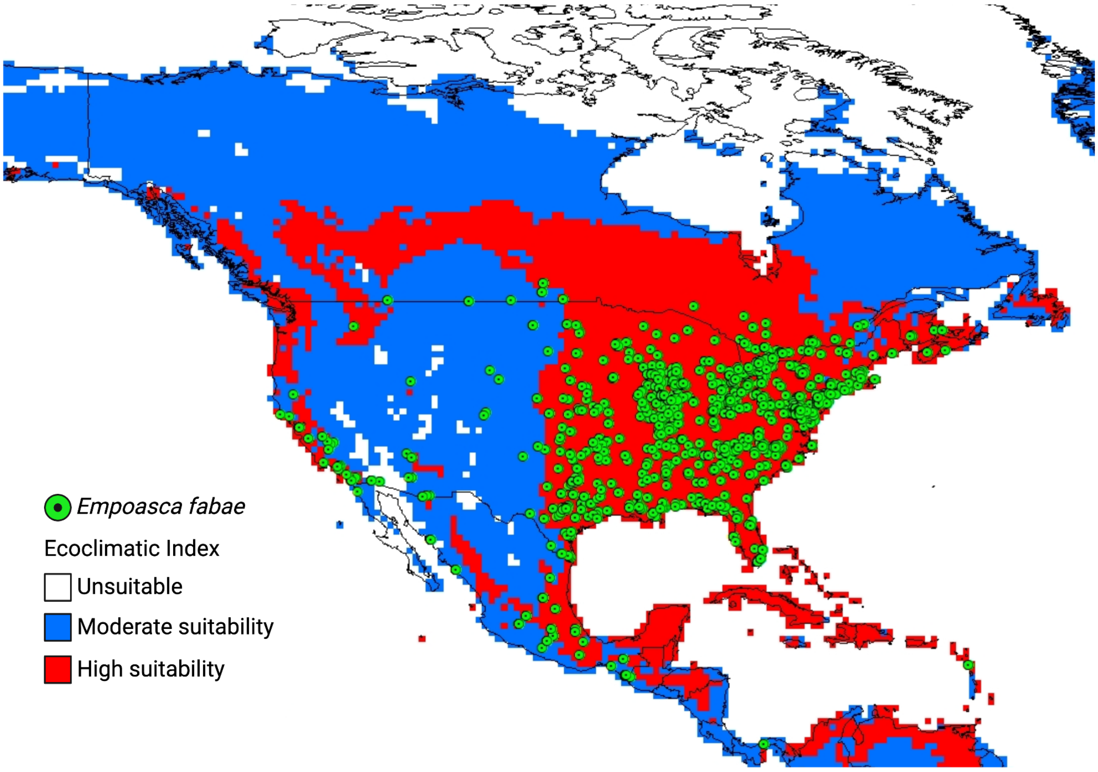


**Supplementary Figure S4.** Potential suitability of North America for E. fabae occurrence based on the Ecoclimatic Index. The green circles indicate areas of occurrence reported on the GBIF database.

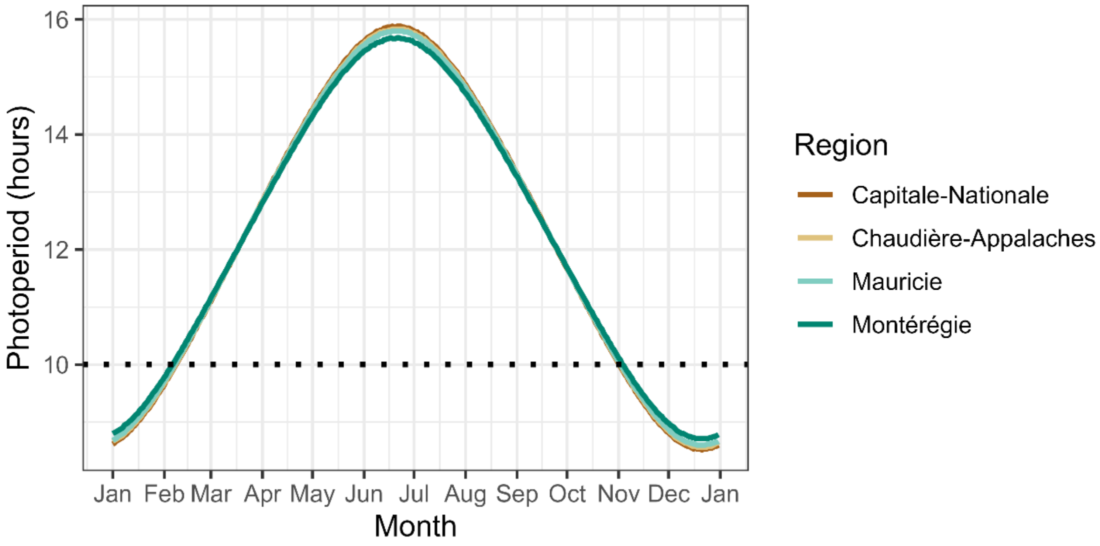


**Supplementary Figure S5**. Photoperiod (hours) in the four geographic regions of Québec (Canada). The dashed line indicates the oviposition threshold where oviposition does not occur (10 hours/day; Taylor et al., 1995).
